# Extracellular ribosomal RNA provides a window into taxon-specific microbial lysis

**DOI:** 10.1101/2021.07.02.450638

**Authors:** Kevin Xu Zhong, Jennifer F. Wirth, Amy M. Chan, Curtis A. Suttle

## Abstract

Microbes are by far the dominant biomass in the world’s oceans and drive biogeochemical cycles that are critical to life on Earth. The composition of marine microbial communities is highly dynamic spatially and temporally, with consequent effects on their functional roles. In part, these changes in composition result from viral lysis, which is taxon-specific and estimated to account for about half of marine microbial mortality. Here we determined taxon-specific cell lysis of prokaryotes in coastal seawater by sequencing extracellular and cellular ribosomal RNA (rRNA). We detected lysis in about 15% of the 16946 prokaryotic amplicon sequence variants (ASVs) identified, and lysis of up to 34% of the ASVs within a water sample. High lysis was most commonly associated with rare but typically highly productive bacteria, while relatively low lysis was more common in taxa that are often abundant, consistent with the proposed model of “kill the winner”, and the idea that less abundant taxa generally experience higher relative lysis than dominant taxa. These results provide an explanation to the long-standing conundrum of why highly productive bacteria that are readily isolated from seawater are often in very low abundance.

**One Sentence Summary:** Extracellular rRNA shows wide variation in cell lysis among prokaryotic taxa.

---

Microorganisms are major drivers of nutrient cycles in the world’s oceans, and constitute more than 90% of its living biomass; yet, have turnover rates ranging from hours to days^1^. A major contributor to these high turnover rates is viral infection that is estimated to kill about 20% of this biomass each day^2^, thereby, shunting cellular organic matter into dissolved carbon and nutrient pools^3-4^. Moreover, viral infection is taxon-specific, contributing to bacterial communities that are highly dynamic in terms of taxonomic composition^5^. Despite the significance of cell lysis to marine ecosystem processes, data on the number and taxonomic composition of bacteria in which lysis occurs remains an enigma.

During investigations into the diversity of RNA viruses in seawater, we were puzzled by the large amount of ribosomal RNA (rRNA) that “contaminated” samples filtered through 0.22-µm pore-size filters. We wondered if the source of the extracellular rRNA (rRNA_ext_) was cell lysis, and if so, could the rRNA_ext_ be sequenced to reveal the taxonomic composition of the cells that had died. The premise of the argument is simple; if the source of the rRNA_ext_ is cell lysis, then sequencing this RNA will reveal the cells from which it was derived, and thus those taxa in which lysis has occurred. Preliminary investigations using quantitative reverse-transcription PCR (qRT-PCR) indicated that there were typically millions of copies of rRNA_ext_ in each mL of 0.22-µm-filtered seawater, and that it was stable for days when incubated on the benchtop. Moreover, it was possible to sequence the rRNA_ext_ and taxonomically profile the community from which it was derived. Here, we detail and expand on these observations to show that rRNA_ext_ is produced by cell lysis of prokaryotes. Moreover, by sequencing and quantifying the rRNA_ext_, cellular rRNA (rRNA_cell_), and the genes encoding ribosomal RNA (rDNA_cell_), we can estimate the taxon-specific cell lysis of prokaryotes in complex natural microbial communities.

## Extracellular rRNA (rRNA_ext_) is produced by cell lysis

We hypothesized that extracellular ribonucleoprotein-bound rRNAs, including free ribosomes, are produced in seawater as the result of cell lysis. To test this hypothesis, we infected the marine heterotrophic proteobacterium, *Vibrio sp*. strain PWH3a, with the lytic phage, PWH3a-P1^6^, and used qRT-PCR to quantify rRNA_ext_ in 0.22-µm filtrate. About 1.75 hours after adding viruses to a culture of *Vibrio sp*. PWH3a, the concentration of rRNA_ext_ was about 100-fold higher than in an uninfected control culture (Fig. 1a), confirming that viral lysis leads to rRNA_ext_ production, likely in the form of ribosomes. Observations by transmission electron microscopy have shown that viral lysis of prokaryotes releases ribosomes^7^. Given that viruses are estimated to kill about 20% of the bacterial standing stock in the oceans each day, viral lysis is likely a significant source for the production of rRNA_ext_.

**Fig. 1.**
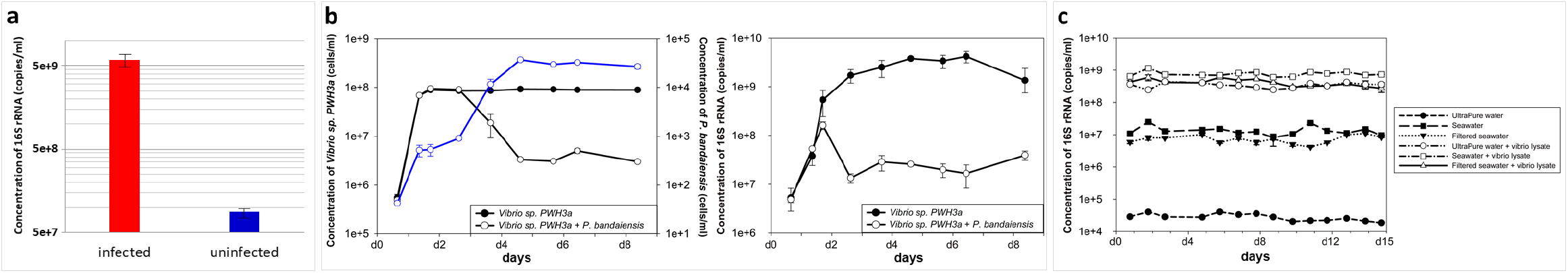
Production and persistence of free extracellular rRNA (rRNA_ext_) in water. **a**: Production of 16S rRNA_ext_ by *Vibrio sp*. strain PWH3a in cultures either infected or not infected with the phage PWH3a-P1 at 1.75 h post-infection. Error bars represent the standard deviation of three replicates. **b:** 16S rRNA_ext_ concentrations in cultures of *Vibrio. sp*. PWH3a, with or without the addition of the grazer, *Paraphysomonas bandaiensis*. The left panel shows changes in the concentration of bacteria (black lines and left axis) and grazer *P. bandaiensis* (blue line and right axis) during eight days of incubation. The right panel shows changes in the concentration of rRNA_ext_, with and without the addition of the grazer. **c:** 16S rRNA_ext_ concentrations in ultrapure water, untreated seawater, and 0.22-µm-filtered seawater, measured over two weeks. Incubations were performed in the dark at 21°C, with and without the addition of 0.22-µm-filtered *Vibrio sp*. PWH3a-P1 lysate (vibrio lysate).

Potentially, grazing by protists could also produce rRNA_ext_. We tested this by adding the bacterivorous flagellate *Paraphysiomonas bandaiensis* to cultures of *Vibrio sp*. PWH3a and quantifying rRNA_ext_. Rather than production, in the presence of grazers, the concentration of rRNA_ext_ decreased relative to that of controls, implying that grazers consume rRNA_ext_ (Fig. 1b). This is consistent with observations that phagotrophic grazers consume virus particles^8^, which are the same size range as ribosomes. Importantly, there was no evidence that grazing of bacteria by protists releases rRNA.

## Extracellular rRNA (rRNA_ext_) is remarkably stable in seawater

The concentration of rRNA_ext_ in seawater is a balance of production and removal; thus, we added rRNA_ext_ produced by lysis of cultures of *Vibrio sp*. PWH3a with phage PWH3a-P1 (Fig. 1c) to molecular grade ultrapure water, untreated seawater and 0.22-µm filtered seawater in order to test its stability. After at least ten days of incubation in the dark at 21°C, there were no detectable changes in the concentrations of rRNA_ext_ in any of the treatments. Collectively, these data demonstrate that the rRNA_ext_ detected by qRT-PCR is remarkably stable in seawater, even in the presence of the natural microbial community. This stability suggests that rRNA_ext_ is in the form of ribosome complexes that prevent degradation of the RNA, consistent with observations that rRNA complexed with ribonucleoproteins is protected from nucleases^9-10^.

Given the stability of rRNA_ext_ in seawater, the taxonomic affiliation of rRNA_ext_ will reflect the microbial community from which the rRNA is derived, provided that the relative abundances are corrected for differences in rRNA_cell_ concentrations that occur among taxa and with physiological state^11^. Therefore, inferring taxon-specific lysis requires dividing the concentration of rRNA_ext_ by rRNA_cell_, for each taxon. Taxon-specific lysis can be compared to estimates of the relative abundance of each taxon, thereby providing an estimate of the relative mortality among taxa. Below, we use this approach to reveal major differences in lysis among microbial taxa in the ocean, and discuss the caveats associated with the method.

## Cell lysis occurs across phyla

We analyzed seawater samples collected from five depths in the Strait of Georgia, British Columbia, Canada, during and after the spring bloom. To infer taxon-specific lysis, we used deep sequencing of a 412-bp amplicon to infer the taxonomic distribution of 16S rRNA_ext_, as well as rRNA_cell_ and rDNA_cell_. A total of 16946 amplicon sequence variants (ASVs), belonging to 33 phyla of bacteria and three phyla of archaea (Fig. 2a) were detected across all ten seawater samples.

**Fig. 2.**
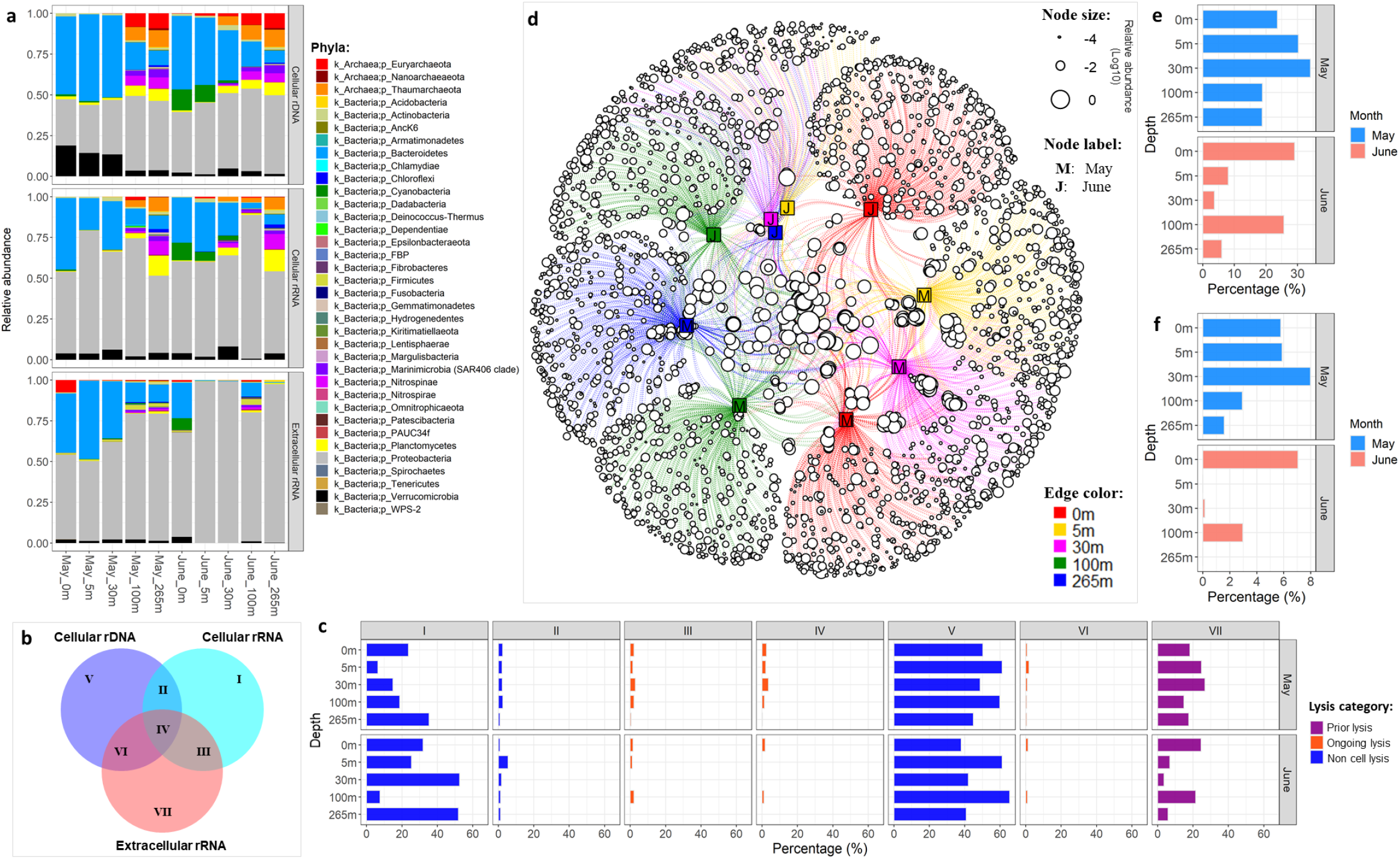
Detection of cell lysis in coastal seawater as inferred from extracellular rRNA (rRNA_ext_). **a**. Relative abundance of 16S amplicons assigned to phyla based on amplicon sequence variants (ASVs) for cellular rDNA and rRNA (rDNA_cell_ and rRNA_cell_, respectively) and rRNA_ext_. **b**. Venn diagram showing seven possible lysis groups, where I, V, and VII represent ASVs found in rDNA_cell_, rRNA_cell_ and rRNA_ext_, respectively; the overlap represents ASVs shared between groups. **c**. Each panel shows the proportion of ASVs (from rDNA_cell_, rRNA_cell_ and rRNA_ext_ pools) across seven lysis groups in each seawater sample. The orange bars represent ASVs with ongoing lysis (group III, IV and VI) as indicated by the presence of both cellular rDNA/rRNA and extracellular rRNA fractions. The purple bars represent ASVs with prior-lysis (group VII) as inferred from the presence of rRNA_ext_ but undetectable rDNA_cell_ or rRNA_cell_. The blue bars indicate ASVs without cell lysis (group I, II and V), as indicated by the absence of rRNA_ext_. **d**. A network showing the taxonomic distribution of ASVs detected in extracellular rRNA, across seawater samples collected from 5 depths. Each open circle (circular node) represents a ribosomal ASV and the size indicates its relative abundance; The line (edge) links ASVs to the seawater samples (square node) where they were detected. It shows that a high proportion of taxa in which lysis was detected were relatively rare and confined to a single month and depth; whereas, there were relatively few taxa in which lysis was detected between months and across depths. **e**. Proportion of all ASVs from each sample (from rDNA_cell_, rRNA_cell_ and rRNA_ext_ pools) in which rRNA_ext_ was detected, indicating cell lysis. **f**. The proportion of ASVs detected as rDNA_cell_ from each sample in which rRNA_ext_ was detected, consistent with ongoing lysis.

For each sample, we binned assigned taxa into seven different “lysis groups” based on their detection as rRNA_cell_, rRNA_ext_ and rDNA_cell_ (Fig. 2b), with their presence as rRNA_ext_ (Groups III, IV, VI, VII) indicating lysis. Across samples, the taxa in which lysis was detected varied widely, ranging from 3.7% to 34.1 % of taxa at 30 m in June and May, respectively (Fig. 2 c-e). Lysis was associated with 28 bacterial and three archaeal phyla, including 19 phyla for which viruses have not been reported (*Acidobacteria, AncK6, Armatimonadetes, Chloroflexi, Epsilonbacteria*, FBP group, *Fibrobacteres, Fusobacteria, Gemmatimonadetes, Hydrogenedentes, Kiritimatiellaeota, Lentisphaerae, Margulisbacteria, Nanoarchaeaeota, Nitrospinae, Planctomycetes*, PAUC34f, *Patescibacteria* and *Spirochaetae*) (Extended Data Fig. 1). Moreover, the proportion of taxa within a phylum in which lysis occurred varied across phyla. In ten of the 31 phyla in which lysis occurred (i.e. *Armatimonadetes, Chlamydiae, Cyanobacteria, Deinococcus-Thermus*, FBP group, *Fibrobacteres, Firmicutes, Fusobacteria, Nanoarchaeaeota, Spirochaetae*), more than half of the ASVs in those phyla were associated with cell lysis, and in some cases included all the taxa within a phylum (Extended Data Fig. 1). Our results show that cell lysis occurred in taxa across a broad range of prokaryotic phyla; yet, for a given sample the maximum percentage of ASVs found in the rRNA_ext_ relative to the rDNA_cell_ (Groups IV and V) was only ∼8% (Fig. 2f), indicating that lysis was not detectable for the vast majority of taxa.

Within taxa in which lysis was detected most were in Group VII (Fig. 2c), indicating that lysis likely occurred prior to sampling, since neither rDNA_cell_ nor rRNA_cell_ was detected. Ongoing lysis, implied by the presence of both rDNA_cell_ and rRNA_ext_ (Groups IV and VI) was detected in fewer taxa. Of these, 0.03% to 1.85% of taxa were detected in the rRNA_cell_, rRNA_ext_ and rDNA_cell_ fractions (Group IV), making it possible to estimate relative mortality, while rDNA_cell_ and rRNA_ext_ (Group VI) occurred in 0.03% to 3.6% of taxa, implying lysis of taxa with little growth or activity. For taxa in Groups I and III, rRNA_cell_ was detected but rDNA_cell_ was undetectable, suggesting they are active but rare, or dormant but harbour high numbers of ribosomes^12^. Up to 2.9% of these taxa were in Group III, with detectable rRNA_cell_ and rRNA_ext_ even though rDNA_cell_ was undetectable, implying that these taxa were in very low abundance but undergoing lysis.

Our observation that lysis was associated with specific ASVs is consistent with viral lysis, which is typically taxon-specific and would be expected to kill cells within a taxon, while leaving members of other taxa unaffected. These observations are congruent with the seed-bank theory of viral infection, in which only a small proportion of viruses are active at any given time^13^. Furthermore, as viral lysis is estimated to cause about 50% of marine prokaryotic mortality^14-15^ and kill about 20% of the standing stock each day^16^, lysis must be high in taxa that are affected. Strikingly, of the 31 phyla for which rRNA_ext_ was detected, there are only a few for which viruses have been isolated (*Euryarchaeota, Thaumarchaeota, Proteobacteria, Firmicutes, Bacteriodetes, Actinobacteria, Cyanobacteria, Chlamydiae, Tenericutes and Deinococcus-Thermus*)^17^ or are known from single amplified genomes (*Verrucomicrobia* and *Marinimicrobia*)^18^. Thus, sequencing of rRNA_ext_ can also provide insights into taxa to be targeted for virus isolation. Although programmed cell death^19^ and predation by *Bdellovibrio* and like organisms (BALOs)^20^ can also cause lysis of prokaryotes, there is no evidence that they are a major source of prokaryotic mortality in the sea.

## Cell lysis varies widely across taxa

Cell lysis can be estimated for individual taxa as the ratio of the relative abundance of rRNA_ext_ to rRNA_cell_; standardizing to cellular rRNA accounts for differences in rRNA concentrations among taxa due to taxonomic or physiological differences. When rDNA_cell_ has not been detected it implies that the cells are in very low abundance or that complete lysis of the population has occurred. When a specific ASV is present in the rRNA_ext_, rRNA_cell_ and rDNA_cell_ pools (Group IV), it implies ongoing lysis of a specific taxon (ASV) that is present in the community (Fig. 3 b-c). Extant populations undergoing lysis were generally few and highly variable across samples; in fact, there were no taxa for which lysis was evident across all samples. Of the 16946 ASVs detected, the number of taxa undergoing lysis (Group IV) ranged between one for the June 30-m sample and 46 for the surface sample in May (Fig. 3b). To gauge whether lysis was high relative to the size of the co-occurring population, we divided rRNA_ext_ by rRNA_cell_ for each ASV. Numbers >1 indicate more free rRNA in the water than in cells for a specific taxon, implying relatively high lysis compared to the size of the population. In our data the values ranged between 0.09 and 5.70, with the highest relative lysis found in members of the phylum *Bacteroidetes*, particularly those in the families *Flavobacteriaceae* and *Saprospiracea* during the phytoplankton bloom in May (Fig. 3C). Copiotrophic bacteria in these families are well known for breaking down high molecular weight dissolved organic material associated with phytoplankton blooms^21^. Similarly, cyanobacteria in the genus *Synechococcus* were relatively abundant, especially at 0 and 5 m in June (Fig. 3a); however, only one of the 138 ASVs assigned to *Synechococcus* was estimated to comprise >1% of the total rDNA_cell_, although we found no evidence of cell lysis for this taxon (Extended Data Fig. 2 and 3). In contrast, during June at 0 and 5 m, lysis was detected in 21 of the less-abundant ASVs assigned to the genus *Synechococcus*, (Fig. 3b; Extended Data Fig. 3). These results are consistent with observations of high host specificity for most cyanophages^22-23^.

**Fig. 3.**
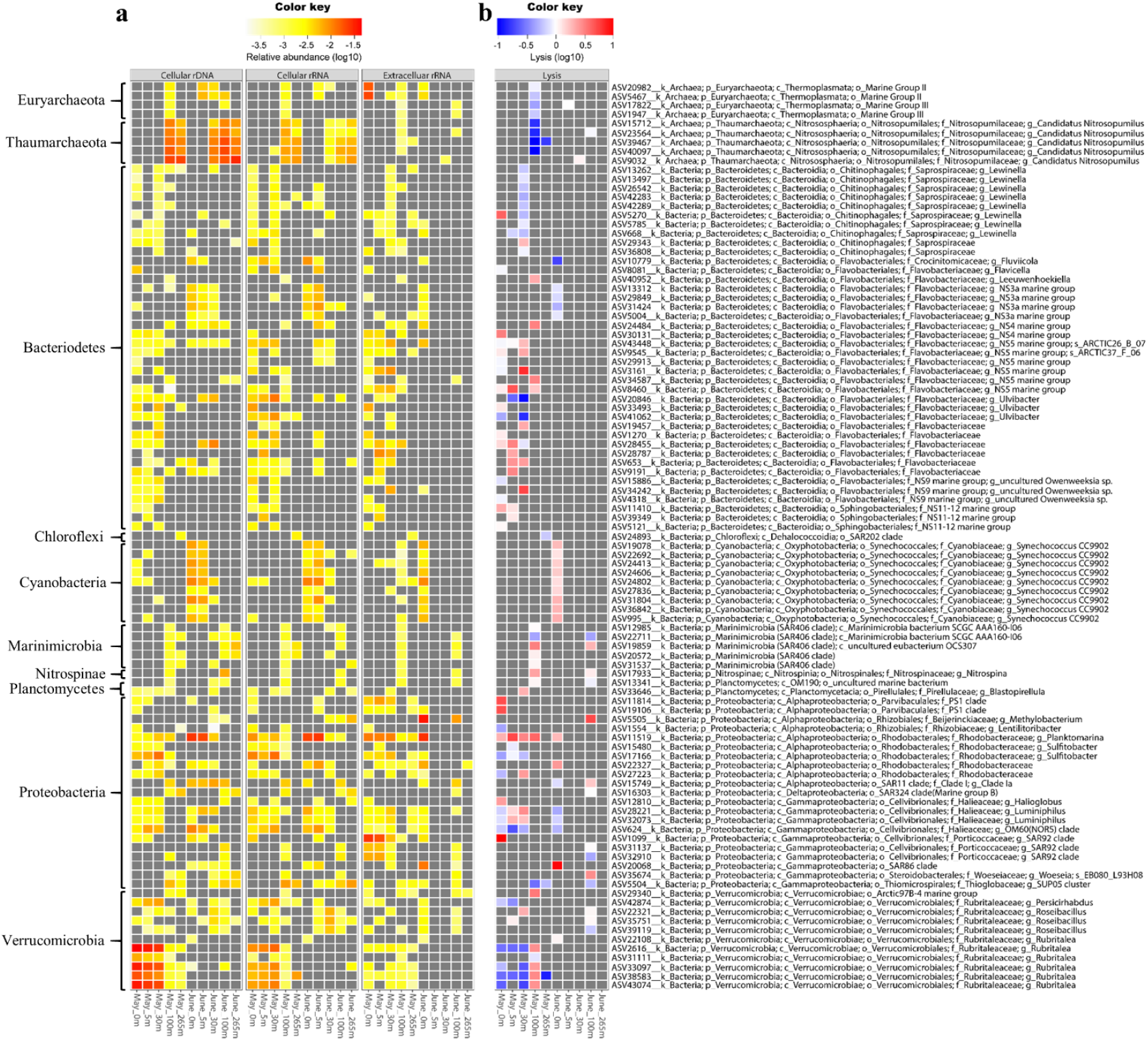
Estimates of taxon-specific cell lysis in coastal seawater samples from the Strait of Georgia. **a**. Panels from left to right show relative abundance of 99 prokaryotic ASVs (Group IV) for rDNA_cell_, rRNA_cell_ and rRNA_ext_. **b**. Taxon-specific lysis measured as the ratio of the relative abundance of rRNA_ext_ to rRNA_cell_, for each of these 99 ASVs in Group IV. In the heatmap, grey indicates that a value required to make the calculation was undetectable.

## High lysis occurs in rare taxa and low lysis occurs in more abundant taxa

An enduring puzzle in marine microbial ecology has been the relationship between viral infection and the structure of prokaryotic communities, and in particular the relationship between microbial abundance and viral lysis for specific taxa. It has been proposed that rates of lysis are relatively low for cells in the most abundant taxa, while relatively rare “r-selected” bacteria that are capable of fast growth account for most of the viral production^24^. This view is congruent with “Kill the Winner”, a model, in which viral lysis prevents the most ecologically “fit” cells from dominating the community^25-26^. Conversely, it has been suggested that cells associated with the most abundant taxa are subject to significant viral lysis^27^. Here, we examined taxa for which we could measure rRNA_ext_, rRNA_cell_ and rDNA_cell_ (Group IV) and demonstrate that high lysis was significantly related to those taxa for which cells were in relatively low abundance (Fig. 4). For example, ASVs assigned to the phylum *Proteobacteria* in the families *Rhodobacteraceae* (genus *Sulfitobacter*) and *Halieaceae* (OM60 clade), and the phylum *Verrucomicrobia* (genus *Rubritalea*) were the most abundant prokaryotes from 0 to 30 m in May (Fig. 3a), and yet estimates of viral lysis were among the lowest measured (Fig. 3b). Similarly, at 0 to 5 m in June, bacteria in the *Rhodobacteraceae* were also among the most abundant bacteria (genus *Planktomarina*), along with members of the phylum *Bacteroidetes* in the family *Flavobacteriaceae* (genus *Formosa*), as well as other ASVs assigned to OM60. Again, although members of these taxa were relatively abundant, lysis was low. Similar observations were made for archaea, which were some of the most abundant cells at 100 m and 265 m, but experienced relatively low lysis (Fig. 3). These included Marine Group I archaea in the phylum *Thaumarchaeota*, which are ubiquitous in the ocean and important in nitrification^28^. Members in the genus of *Candidatus* Nitrosopumilus, which oxidizes ammonia to nitrite, were estimated to account for up to 12.9% of the 16S sequences below 30 m in May, and up to 14.5% of the sequences in June, but showed consistently low lysis. Another example is proteobacteria in the order *Pelagibacterales*, (*a*.*k*.*a*. SAR11 clade), which are among the most abundant bacteria in the open ocean. There were 612 ASVs assigned to SAR11, which encompassed three putative families (Clades I, II, and III). Members of Clade I were relatively abundant in our samples from 100 m and 265 m in May and at all depths in June (Extended Data Fig. 2); yet, with the exception of one ASV at 100 m in June (Fig. 3), extracellular rRNA from the SAR11 group was undetectable, indicating negligible cell lysis. Even though bacteria in the SAR11 clade were relatively abundant in several samples, and are known to be infected by a variety of viruses^27^, we could detect little to no lysis (Extended Data Fig. 2). Collectively, these data indicate that much of the cell lysis and release of rRNA that occurs in seawater can be attributed to prokaryotic cells that are typically present in relatively low abundance, while populations of cells in relatively high abundance typically have low rates of cell lysis.

**Fig. 4.**
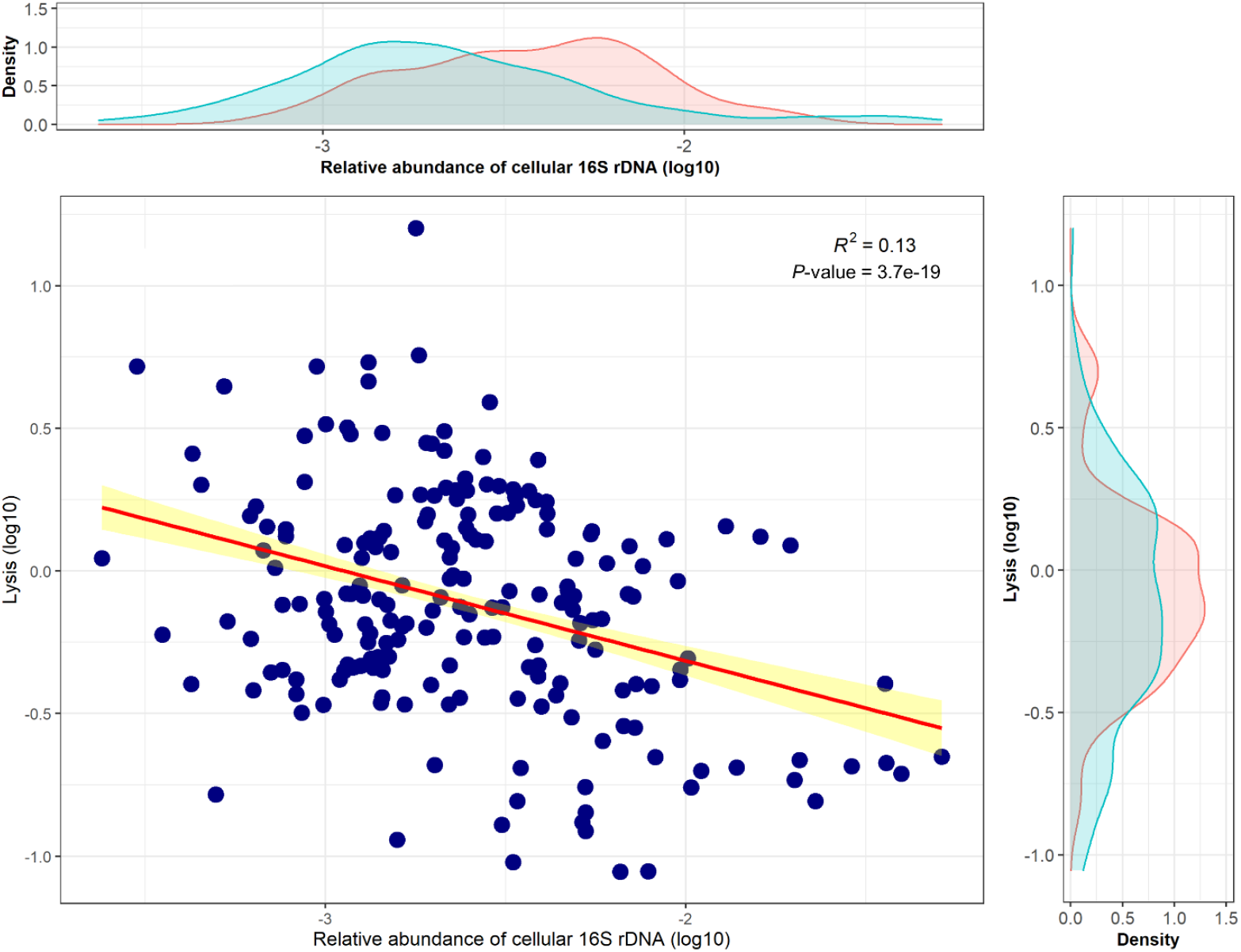
Linear regression (red line) between the relative abundance of 16S rDNA_cell_and lysis for individual ASVs across all samples. These represent the 99 ASVs that belong to Group IV in which rDNA_cell_, rRNA_cell_ and rRNA_ext_ were detectable, allowing relative lysis to be estimated. Yellow color represents the confidence interval of regression coefficients. The density plots above and to the right of the data show the distribution of values for May (pink) and June (green).

## Turnover of extracellular RNA (rRNA_ext_)

We found that rRNA_ext_, is stable when incubated for two weeks in seawater in the dark at 21°C, even in the presence of the natural microbial community (Fig. 1c). However, in natural waters other factors also contribute to decay of rRNA_ext_. For example, solar radiation causes photodegradation of organic molecules in seawater^29^, and increases decay rates of viral infectivity by more than ten-fold^6,30^, and would be expected to have similar effects on the degradation of rRNA_ext_. Furthermore, large differences in the taxonomic profiles of rRNA_ext_ across depths and between sampling times, indicates that the taxonomic composition of the ASVs in seawater is dynamic. Moreover, there is no *a priori* reason to expect rRNA_ext_ from different taxa to decay at different rates, because the potential decay mechanisms (e.g. solar radiation, temperature, protistan grazing) are unlikely to be taxon-specific. Thus, the turnover rate of rRNA_ext_ will not affect calculations of relative taxon-specific lysis, which is the ratio of the relative abundance of rRNA_ext_ to rRNA_cell_, and provides an estimate of relative mortality among taxa. Hence, the conclusion that high lysis is coupled with low abundance is not affected by the turnover rate of rRNA_ext_.

The determination of absolute lysis rates of different taxa requires estimates of turnover rates of rRNA_ext_. Although we do not have absolute estimates of turnover rates, ultimately rRNA_ext_ decay must be balanced by production. For example, if the decay and production rates of rRNA_ext_ is 1 d^-1^, which is also a typical bacterial production rate for coastal seawater, then the taxon-specific lysis rate will be equal to the ratio of the concentration of rRNA_ext_ to rRNA_cell_ for each taxon multiplied by the rRNA_ext_ turnover rate (Extended Data Fig. 4).

In summary, we have shown that extracellular RNA (rRNA_ext_) in seawater is produced by cell lysis, and not by protistan grazing on bacteria. Although our approach cannot distinguish among different causes of cell lysis, given that lysis by viruses accounts for about half of the mortality of prokaryotes in seawater^14-15^, and kills about 20% of bacteria each day^16^, it is reasonable to infer that most rRNA_ext_ stems from viral infection. Moreover, rRNA_ext_ is stable in seawater and can be sequenced to taxonomically profile the cells in which lysis has occurred. We demonstrate that there is widespread lysis across prokaryotic phyla, but that within a sample, lysis is only detected in a low proportion of taxa. Relative taxon-specific lysis, which is relative to the taxon-specific mortality rate, could be measured using the ratio of the relative abundance of rRNA_ext_ to rRNA_cell_. Our results indicate that high lysis is significantly related to low abundance, suggesting that, overall, rare taxa are associated with high lysis; conversely, low lysis is associated with high abundance, suggesting that the dominant taxa are subject to low relative rates of lysis. Thus, the ability to estimate taxon-specific lysis adds an important tool in our quest to explain the distribution and abundance of specific microbial taxa in nature.

## Data Availability

Sequencing data generated in this study have been deposited in the NCBI Sequence Read Archive (SRA) under the accession numbers SRR14873150 to SRR14873179.

## Code Availability

The related codes for analyzing the taxon-specific lysis are included in the custom R package: tslysis (https://github.com/kevinzhongxu/tslysis).

## Acknowledgments

We thank members of the Hakai Institute for facilitating the collection of seawater samples, particularly Brian Hunt, Kate Lansley, Alex Hare and Megan Foss. We are grateful to David Caron for providing *Paraphysiomonas bandaiensis* for the grazing studies. This work was supported by grants to CAS from the Tula Foundation, the Gordon and Betty Moore Foundation, a Discovery grant from the Natural Sciences and Engineering Research Council of Canada, and infrastructure awards from the Canada Foundation for Innovation and the British Columbia Knowledge Development Fund.

## Author Contributions

KXZ designed experimental approaches, conducted experimental work, analyzed the data, wrote the initial draft of the manuscript, and oversaw subsequent versions. JFW conducted initial experimental work and edited the manuscript. AMC provided technical support throughout the project and edited the manuscript. CAS conceived the project, contributed to experimental design and data interpretation, and helped write the paper.

## Competing interests

The authors declare no competing interests.

## Supplementary Materials

### Methods

#### Culturing conditions

Heterotrophic marine bacterium *Vibrio sp*. strain PWH3a and its phage PWH3a-P1^6^ were grown in CPM medium (0.05% Casamino Acids [Difco], 0.05% Bacto Peptone [Difco] in ultrafiltered ∼25 psu seawater)^6^. A culture of the phagotrophic flagellate *Paraphysomonas bandaiensis* (courtesy David Caron, University of Southern California) was grown with its associated bacterial community in F/2-enriched seawater^31^ to which a rice grain was added^32^. All media were autoclave-sterilized; cultures were maintained at room temperature (∼21°C) in the dark.

#### Seawater sampling and filtration

Seawater samples were collected from Hakai Oceanographic station QU#39 (50.0307N, 125.0992W) in the Strait of Georgia near Quadra Island, British Columbia, Canada, during (May 5, 2015) and after (June 4, 2015) a spring bloom. One-liter seawater samples were collected from five depths (0, 5, 30, 100 and 265 m) using Niskin bottles.

Filtration was used to separate extracellular rRNA (rRNA_ext_) from the cells. Briefly, 100 mL of seawater was pre-filtered through a 120-μm mesh-size Nitex screen to remove large particles, followed by gentle vacuum filtration through a 47-mm 0.22-µm pore-size PVDF filter (Millipore, GVWP). The rRNA_ext_ was collected in the <0.22-µm filtrate, and the cellular rRNA (rRNA_cell_) was retained on the filter. Both the filter and filtrate were flash-frozen in liquid nitrogen and kept at −80°C prior to nucleic acids extraction. The filtrate was divided into several aliquots prior to freezing, so that subsamples could be thawed for downstream applications. The absence of cells in the filtrate was verified by flow cytometry (FCM) and quantitative PCR (qPCR) using primers that target the 16S rRNA gene. See below for further details.

#### Assay to examine rRNA_ext_ production by bacteria infected with viruses

To assess whether viral lysis resulted in rRNA_ext_ production, *Vibrio. sp*. PWH3a was cultured alone (control) and with its phage PWH3a-P1. Briefly, exponentially growing *Vibrio. sp*. PWH3a cells were inoculated into 50 mL of CPM medium in a 250-mL Erlenmeyer flask. After 1.75 h the cells were split into two cultures; one was infected with phage PWH3a-P1 at a multiplicity of infection of ∼10, while the other served as an untreated control. Approximately 1.75 hours after infection, the cultures were filtered through 0.22-µm pore-size syringe filters (Durapore®, PVDF, 33 mm diameter, Millipore) to collect the filtrate containing the rRNA_ex_. Filtrate samples were flash-frozen in liquid nitrogen and stored at −80°C until the rRNA_ex_ was quantified in triplicate samples by reverse-transcription qPCR (RT-qPCR).

#### Assay to examine rRNA_ext_ production by protist grazing on bacteria

To determine if grazing by protozoa resulted in rRNA_ext_ production, we added exponentially growing *Vibrio sp*. PWH3a to a culture of the phagotrophic flagellate *P. bandaiensis* growing in 100 mL of 20% CPM medium (bacteria:grazers = 12500:1 at T0). *Vibrio. sp*. PWH3a grown in the absence of the grazer was used as a control. The experiment was conducted in triplicate in the dark at room temperature (21°C) in 250 mL polycarbonate Erlenmeyer flasks (Corning) for nine days. The cultures were subsampled (250 µL) once or twice a day to count the bacteria and protists by flow cytometry (FCM). Another 1 mL was taken from each flask and filtered through a 0.22-µm pore-size syringe filter (Durapore®, PVDF, 33 mm diameter, Millipore), and the filtrate flash-frozen in liquid nitrogen and stored at −80°C until the concentration of rRNA_ext_ was determined by qRT-PCR.

#### Stability of rRNA_ext_ in water

The stability of rRNAext in water was assayed by adding 100 µL of 0.22-μm-filtered PWH3a-P1 lysate of *Vibrio sp*. PWH3a to 35 mL of Ultrapure water (Invitrogen), unfiltered seawater or 0.22-μm-filtered seawater. Each treatment was conducted in duplicate and incubated in the dark at room temperature (21°C) in 50 mL polypropylene tubes (BD Falcon). Every day for two weeks, 500 µL was taken from each tube, 0.22-µm filtered, flash frozen in liquid nitrogen and stored at −80°C until the concentration of rRNA_ext_ was determined by qRT-qPCR. All filtrations were done using syringe filters (Durapore®, PVDF, 33 mm diameter, Millipore).

#### Flow cytometry (FCM) to count bacteria, viruses and protists

Bacteria, virus and protist numbers were determined using a FACSCalibur flow cytometer (Becton Dickinson) equipped with an air-cooled laser (15 mW, 488 nm) following established methods^33-34^. The flow cytometer list mode files were analysed using WEASEL v3.1 (The Walter and Eliza Hall Institute of Medical Research).

Samples of bacteria and viruses were fixed with EM-grade glutaraldehyde (0.5% final concentration) for 15 min in the dark at 4°C, then flash frozen in liquid nitrogen and stored at - 80°C until analysis. Prior to analysis, samples were diluted in 0.02-µm-filtered TE buffer (10 mM Tris-HCL and 1 mM EDTA, pH 8), and stained with SYBR Green I (at a final 5 × 10^−5^ dilution of the commercial stock solution, Molecular Probes), for 15 min at room temperature (21°C) for bacteria counts, or at 80°C for virus counts.

Protist numbers were determined after staining live cells with LysoTracker Green® (Molecular Probes)^35^. Briefly, flagellates were stained by adding 25 µl of 1 mM dye to 250 µl of culture and incubated for 15 min in the dark at room temperature.

#### Nucleic acids extraction

Total (cellular) nucleic acids (DNA and RNA) were extracted from the 0.22-µm pore-size filters using a MasterPure™ Complete DNA and RNA Purification Kit (Epicentre) by following the manufacturer’s directions for tissue samples. Half of each total nucleic acid sample was treated with 1µL RNase A (Epicentre) at 37°C for 30 min to obtain cellular DNA; the other half of the sample was treated with DNase I (Amplification Grade, Invitrogen) to obtain cellular RNA. Briefly, the DNase I treatment was conducted at room temperature (21°C) for 15 min in a 20 µL reaction mixture that included 16 µL of total nucleic acids (<2 µg) and 2 µL of DNase I (1 U/µL). The reaction was stopped by adding 2 µL of 25 mM EDTA solution and heating at 65°C for 10 min. The purified cellular RNA was immediately used for cDNA synthesis step.

Extracellular nucleic acids (DNA and RNA) were extracted from 400 µL of the <0.22-µm filtrate using a PureLink Viral RNA/DNA mini Kit (Invitrogen) by following manufacturer’s directions and eluted in 25 µL molecular grade water. To obtain the RNA fraction only, DNA was removed by DNase I (Amplification Grade, Invitrogen) treatment by following the manufacturer’s instruction as shown above. The removal of DNA was verified by qPCR to quantify the 16S rRNA gene before and after DNase I treatment. The purified extracellular RNA was immediately applied to the following cDNA synthesis.

#### cDNA synthesis

The DNA-free cellular and extracellular RNA samples were subsequently reverse-transcribed to the complementary DNA (cDNA) by using SuperScript™ III Reverse Transcriptase (Invitrogen) following the manufacturer’s directions. The cDNA samples were then used as the template in quantitative PCR (qPCR) to estimate rRNA copies, and in PCR for deep sequencing of amplicons to obtain the taxonomic distribution of cellular and extracellular rRNAs.

#### Quantitative PCR (qPCR)

Quantitative PCR, using primer set 331F/518R (Extended Data Table 1) to target conservative V3 region of 16S rRNA gene, was carried out on the following 3 fractions:

1. **cellular DNA** obtained from the 0.22-µm pore-size filter to estimate the number of 16S rRNA genes (rDNA_cell_),
2. **cDNA of cellular RNA** to estimate the concentration of cellular 16S rRNA (rRNA_cell_) using cDNA from the 0.22-µm pore-size filter
3. **extracellular RNA** to estimate the concentration of extracellular 16S rRNA (rRNA_ext_) using cDNA from the <0.22-µm filtrate.

In this study, when qPCR was applied to cDNA, it is referred to as quantitative reverse-transcription PCR (qRT-PCR).

Briefly, the 10 µL qPCR reaction contained 1 X SsoFast™ EvaGreen® Supermix (Bio-Rad), 0.5 µM of each primer, and 1 µL of cDNA or DNA template. Thermal cycling was conducted in a CFX96 real-time PCR detection system (Bio-Rad) with the following program: 3 min denaturation at 95°C, followed by 40 cycles of denaturation at 95°C for 30 s, annealing and extension at 62.8°C for 30 s. Nine 10-fold serially diluted standards (ranging from 5 × 10 to 5 × 10^9^ molecules per mL) were run in duplicate along with two no-template control reactions containing 1 µL of nuclease-free water. The amplicon standards were made from a cloned 16S rRNA gene amplified from *Vibrio sp*. PWH3a using primer set 331F/518R, purified using a MiniElute® PCR Purification Kit (Qiagen), and quantified using a Qubit® dsDNA High Sensitivity Assay Kit (Invitrogen). The size of the amplicon (i.e. 187 bp) was verified using gel-electrophoresis, and the qPCR melting curves confirmed that the fluorescence signal corresponded to a DNA fragment of a single size. The qPCR amplification efficiency was between 0.95 and 1.05 for the cloned amplicons (r>0.98, n=9).

#### PCR, amplicon sequencing library construction and sequencing

PCR was conducted to amplify the 16S rRNA gene sequences from DNA or cDNA among samples of each fraction, including cellular DNA and cDNA from the 0.22-µm pore-size filter, and extracellular cDNA from <0.22-µm filtrate, to examine the taxonomic distribution of 16S rDNA_cell_, rRNA_cell_ and rRNA_ext_, respectively, using amplicon deep sequencing.

The preparation of 16S amplicon libraries was adapted from the online Illumina protocol^39^ with several modifications. Briefly, two successive runs of PCR were performed. The first PCR generated amplicons of the 412-bp 16S rRNA gene between the V4 and V5 regions using the modified primers 515F-Nxt and 926R-Nxt (Extended Data Table 1). Compared to 515F and 926R^38^, the modified primers incorporate overhanging adapter sequences (Extended Data Table 1) that are compatible with Illumina index and sequencing adapters. These modifications enabled the use of Illumina Nextera XT indexes as forward and reverse primers to create the dual-indexed amplicon libraries in the second PCR.

##### First amplicon PCR

The 25 μL reaction mix consisted of 1X PCR buffer, 4 mM MgCl_2_, 50 µg of Bovine Serum Albumin (Invitrogen), 200 mM of each dNTP (Invitrogen), 0.4 µM of each primer, 0.5 U of Q5® high fidelity polymerase (NEB), and approximately 5 ng of DNA or 0.5 ng of cDNA template. Each sample was amplified in triplicate.

The PCR program was as described in ref. 38; 25 cycles were used for rDNA_cell_ and rRNA_cell_ samples but the number of cycles was increased to 34 for the rRNA_ext_ samples. Briefly, PCR reactions were subject to an initial denaturation at 95°C for 3 min, followed by 25-34 cycles of denaturation at 95°C for 45 s, annealing at 50°C for 45 s, and elongation at 68°C for 90 s, and a final elongation step at 68°C for 5 min to ensure complete amplification. Triplicate first PCR products were pooled and then purified using magnetic Agencourt AMPure XP beads (Beckman Coulter) with a ratio of 1:1 for beads:product to remove fragments less than 200bp (e.g. dimers). Purified amplicons from the first PCR were then used as the template for the index-PCR (second PCR).

##### Second PCR (index-PCR)

PCR reactions followed the Illumina protocol^39^, but substituting Q5® high fidelity polymerase (NEB). The 25 μL reaction mix consisted of 1X PCR buffer, 4 mM MgCl_2_, 200 mM of each dNTP (Invitrogen), 2.5 µL of each index primer (N7XX and S5XX of Nextera® XT Index Kit), 1 U of Q5® high-fidelity polymerase (NEB) and 2.5 µL of purified DNA product from the first PCR.

Second PCR reactions were subject to an initial denaturation at 95°C for 3 min, followed by 12-16 cycles of denaturation at 95°C for 30 s, annealing at 55°C for 30 s and elongation at 72°C for 30 s, and a final elongation step for at 72°C for 10 min. Twelve cycles for rDNA_cell_ and rRNA_cell_, and 16 cycles (for rRNA_ext_) of index-PCR were used to append Illumina Nextera XT indexes to each side of the custom-designed amplicons.

Cleanup was conducted using magnetic Agencourt AMPure XP beads (Beckman Coulter) to purify amplicons >200bp with a ratio of 1:1 for beads:product. Amplicon libraries were quantified using a Qubit® dsDNA HS Assay Kit (Invitrogen), and the average fragment size was determined using an Agilent Bioanalyser and Agilent High Sensitivity DNA Kit. Equimolar amounts of purified amplicons from the second PCR were pooled for each library. The multiplexed pool was sequenced at the UCLA Sequencing and Genotyping Core facility, using Illumina MiSeq 2 × 300bp paired-end chemistry.

##### Sequence analysis

Amplicon sequences from each of the rDNA_cell_, rRNA_cell_ and rRNA_ext_ fractions were processed and analyzed using the QIIME pipeline version 2 (qiime2.2018.06)^40^. Briefly, adapters and low-quality reads were trimmed with Trimmomatic-0.36^41^ using the following settings (EADING:3 TRAILING:3 SLIDINGWINDOW:4:15 MINLEN:36). Paired-end reads were then merged using PEAR^42^ with the default setting. On average, 204970 ± 95344 merged reads of ∼412 bp were obtained per library; and only about 3% low-quality reads needed to be discarded. These assembled sequences were then loaded into the QIIME pipeline version 2 and DaDa2^43^ was used to filter noisy and chimeric sequences to obtain amplicon sequence variant (ASV) features. Taxonomy was assigned for these ASV features using a pre-built Naïve Bayes classifier that was trained based on the Silva v123 16S SSU database^44^ at 99% nucleotide sequence similarity.

ASV features were removed for downstream analysis for those with taxonomy-assignment confidence less than 80%, with no more than ten reads in either sample, unclassified at the kingdom level, or belonging to eukaryotes, mitochondria and chloroplasts. In the end, for each fraction (rDNA_cell_, rRNA_cell_ and rRNA_ext_), samples were normalized by analyzing the relative abundance for each ASV as the proportion of all sequences within a sample. Statistical analysis was conducted in R version 4.0.3^45^ and figures were generated using ggplot2 version 3.3.3^46^. The network showing the taxonomic distribution of rRNA_ext_ ASVs across seawater samples was plotted using R package igraph v1.2.6^47^. Linear regression analysis to reveal the relative abundance of cellular rDNA and lysis was conducted using R package methods v3.6.2^45^, ggplot2 and ggpmisc v0.3.7^48^.

#### Taxon-specific lysis calculation

Estimate of taxon-specific lysis and identification of lysis groups were performed using the custom R package: tslysis (https://github.com/kevinzhongxu/tslysis). Briefly, for a taxon (ASV) of a given sample, we calculated lysis using the ratio of relative abundance of rRNA_ext_ to rRNA_cell_, as follows.

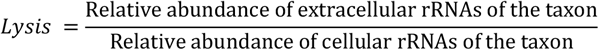

**Extended Data Table 1.**
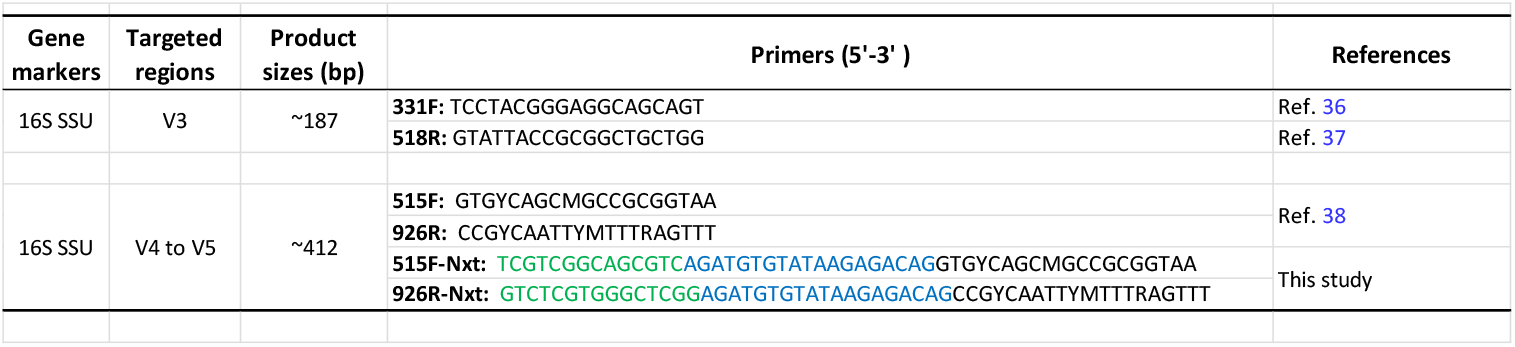
List of primers used in this study.

**Extended Data Fig. 1.**
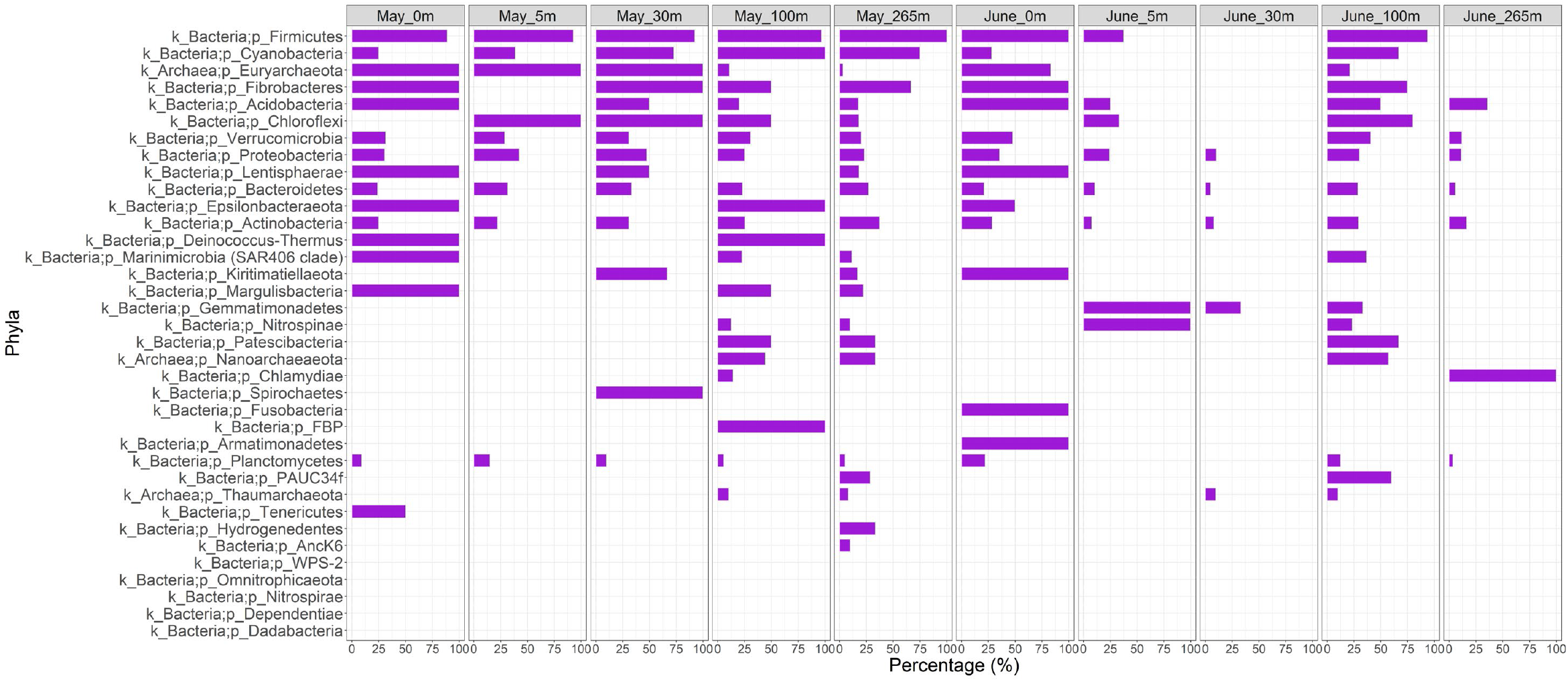
Proportion of ASVs in each sample (from 16S rDNA_cell_, rRNA_cell_ and rRNA_ext_) in which extracellular rRNA was detected, indicating cell lysis.

**Extended Data Fig. 2.**
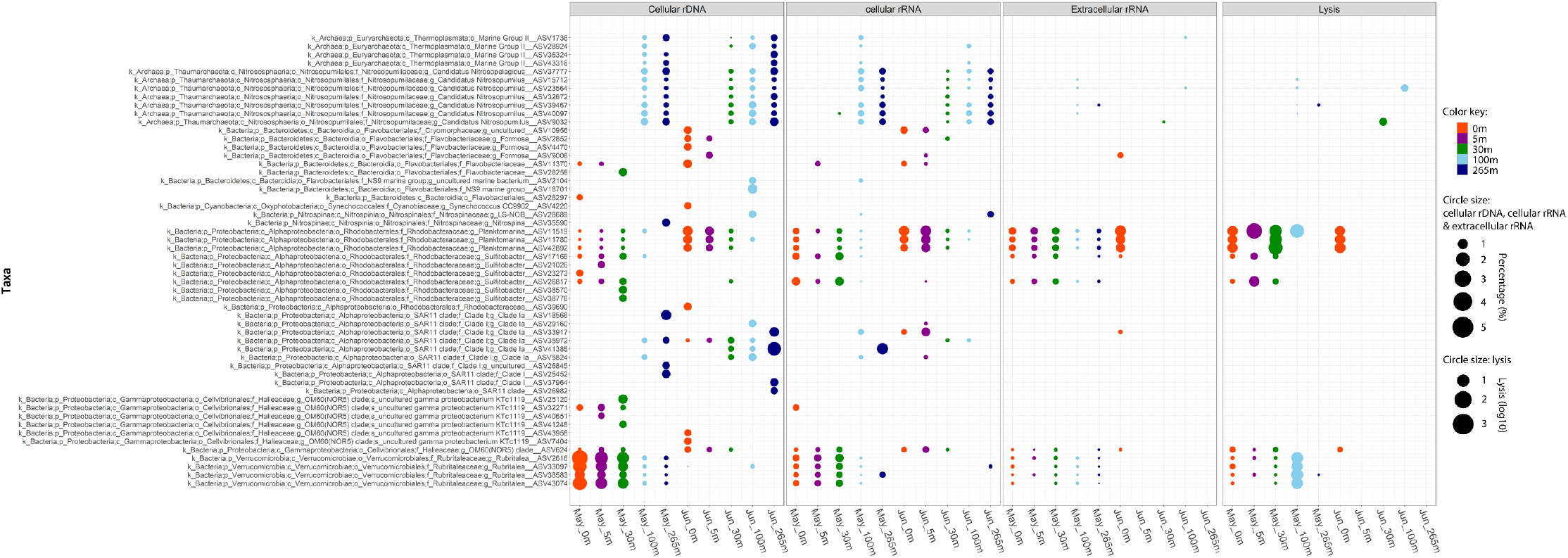
Relative abundance of 16S rDNA_cell_, rRNA_cell_ and rRNA_ext_, and taxon-specific lysis for the dominant prokaryotic ASVs (relative abundance of cellular rDNA >1%) in coastal seawater samples from Strait of Georgia. Taxon-specific lysis was measured as the ratio of the relative abundance of rRNA_ext_ to rRNA_cell_ for each ASV.

**Extended Data Fig. 3.**
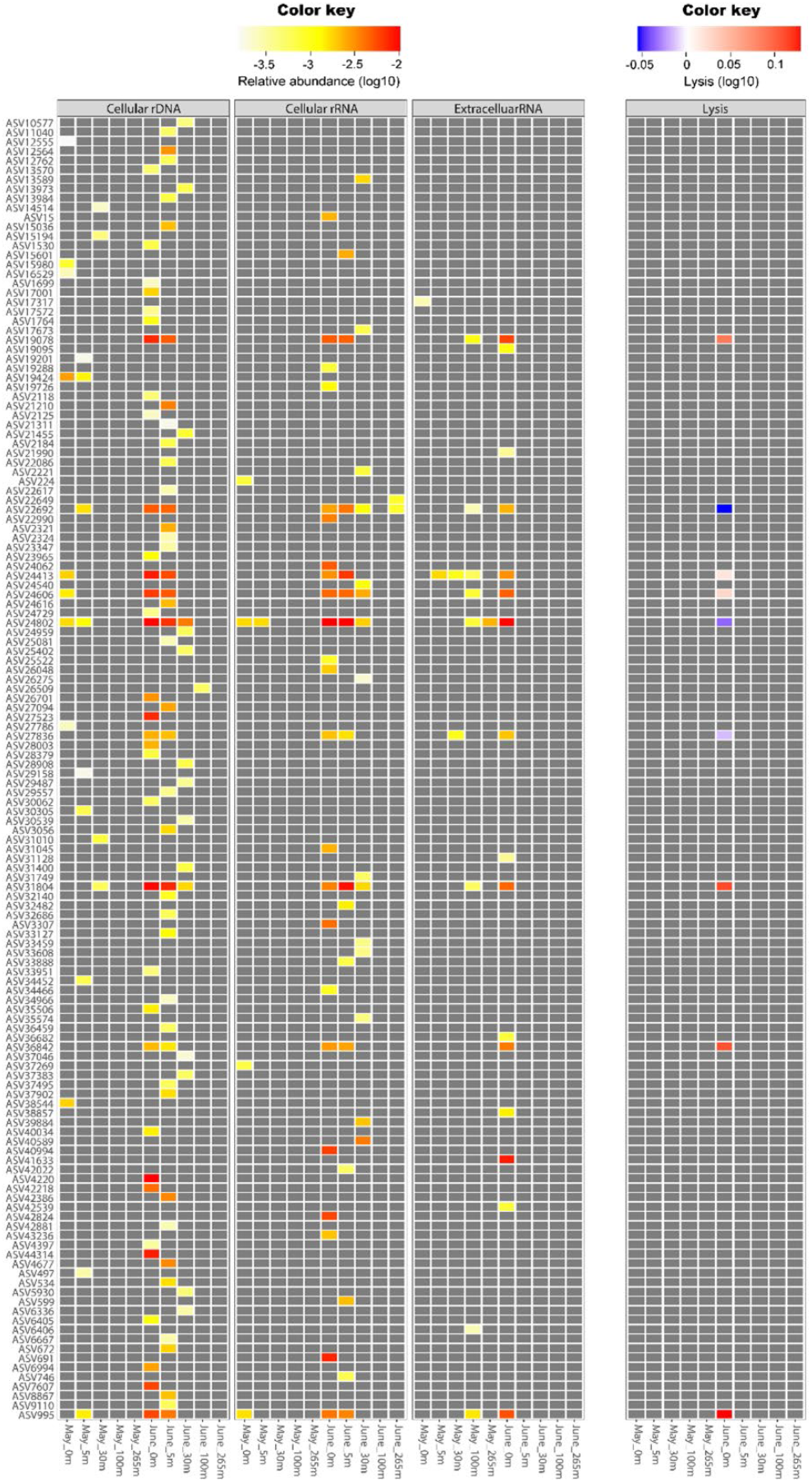
Relative abundance of 16S rDNA_cell_, rRNA_cell_ and rRNA_ext_, and taxon-specific lysis for 138 *Synechococcus* ASVs obtained from ten seawater samples from Strait of Georgia. Taxon-specific lysis was measured as the ratio of the relative abundance of rRNA_ext_ to rRNA_cell_ for each ASV. In the heatmap, grey indicates that a value required to make the calculation was undetectable.

**Extended Data Fig. 4.**
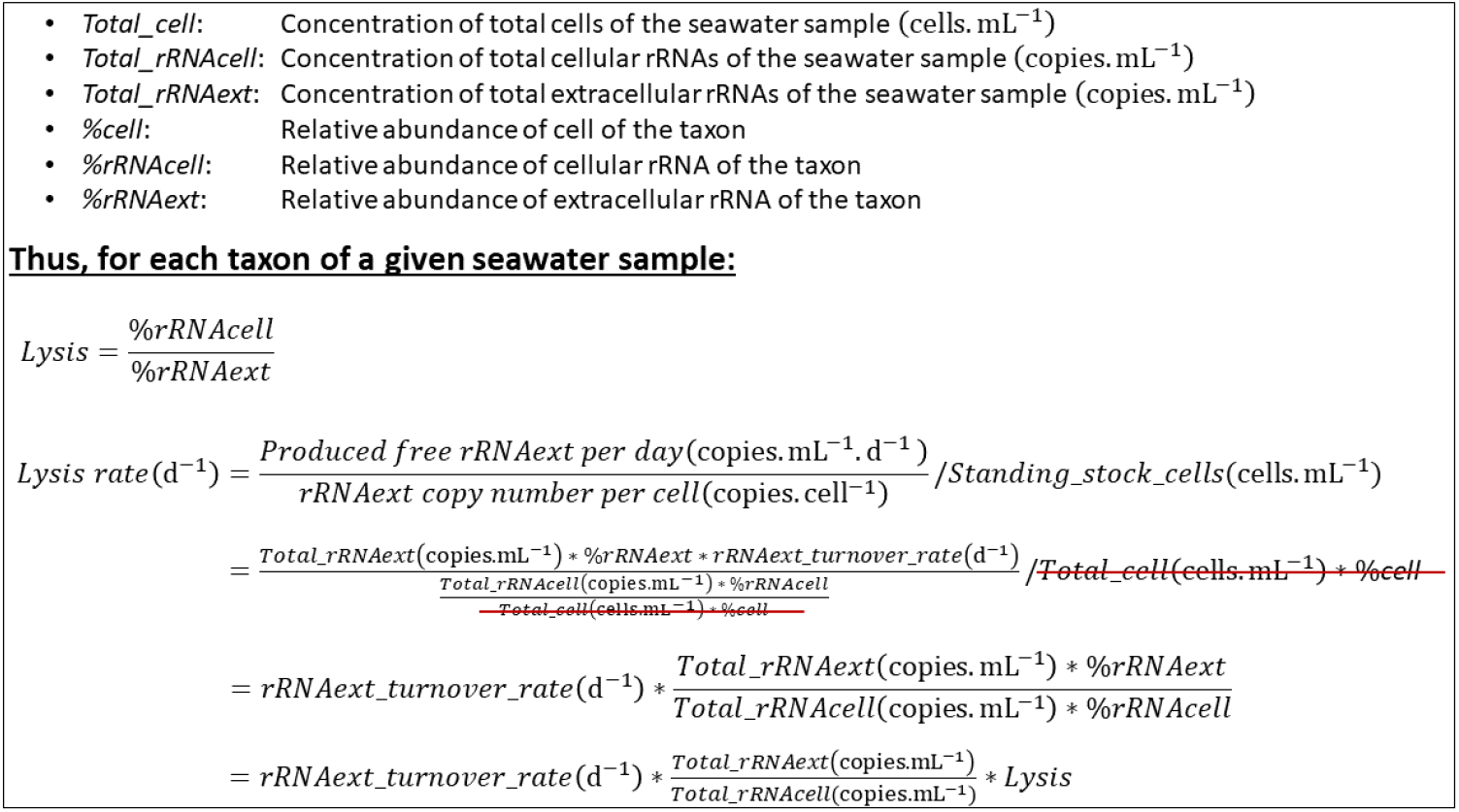
Calculation of the taxon-specific lysis rate, and its relationship with the taxon-specific lysis.

## Notes

https://github.com/kevinzhongxu/tslysis

### Competing Interest Statement

The authors have declared no competing interest.

## References

1. Fuhrman, J. A., Cram, J. A. & Needham, D. M. Marine microbial community dynamics and their ecological interpretation. Nat. Rev. Microbiol. 13, 133–146 (2015).

2. Suttle, C. A. Viruses in the sea. Nature 437, 356–361 (2005).

3. Wilhelm, S. W. & Suttle, C. A. Viruses and nutrient cycles in the sea: Viruses play critical roles in the structure and function of aquatic food webs. BioScience. 49, 781–788 (1999).

4. Fuhrman, J. A. Marine viruses and their biogeochemical and ecological effects. Nature 399, 541–548 (1999).

5. Weinbauer, M. G. Ecology of prokaryotic viruses. FEMS Microbiol. Rev. 28, 127–181 (2004).

6. Suttle, C. A. & Chen, F. Mechanisms and rates of decay of marine viruses in seawater. Appl. Environ. Microbiol. 58, 3721–3729 (1992).

7. Dai, W. et al. Visualizing virus assembly intermediates inside marine cyanobacteria. Nature 502, 707– 710 (2013).

8. González, J. & Suttle, C. Grazing by marine nanoflagellates on viruses and virus-sized particles: ingestion and digestion. Mar Ecol Progr. Ser. 94, 1–10 (1993).

9. Datta, A. K. & Burma, D. P. Association of ribonuclease I with ribosomes and their subunits. J. Biol. Chem. 247, 6795–6801 (1972).

10. Deutscher, M. P. Maturation and degradation of ribosomal RNA in bacteria. Prog. Mol. Biol. Transl. Sci. 85, 369–391 (2009).

11. Bremer, H. & Dennis, P. P. in Neidhardt et al., Eds. (American Society for Microbiology, 2nd ed., 1996) chap. 97, pp. 1559, Tab. 3.

12. Blazewicz, S. J., Barnard, R. L., Daly, R. A. & Firestone, M. K. Evaluating rRNA as an indicator of microbial activity in environmental communities: limitations and uses. ISME J. 7, 2061–2068 (2013).

13. Breitbart, M. & Rohwer, F. Here a virus, there a virus, everywhere the same virus? Trends Microbiol. 13, 278–284 (2005).

14. Fuhrman, J. A. & Noble, R. T. Viruses and protists cause similar bacterial mortality in coastal seawater. Limnol. Oceanogr. 40, 1236–1242 (1995).

15. Mojica, K. D. A. & Brussaard, C. P. D. Significance of viral activity for regulating heterotrophic prokaryote community dynamics along a meridional gradient of stratification in the Northeast Atlantic Ocean. Viruses 12, 1293 (2020).

16. Suttle, C. A. The significance of viruses to mortality in aquatic microbial communities. Microb. Ecol. 28, 237–243 (1994).

17. Bibby, K. Improved bacteriophage genome data is necessary for integrating viral and bacterial ecology. Microb. Ecol. 67, 242–244 (2014).

18. Labonté, J. M. et al. Single-cell genomics-based analysis of virus–host interactions in marine surface bacterioplankton. ISME J. 9, 2386–2399 (2015).

19. Bayles, K. W. Bacterial programmed cell death: making sense of a paradox. Nat Rev Microbiol. 12, 63– 69 (2014).

20. Sockett, R. E. Predatory lifestyle of Bdellovibrio bacteriovorus. Annu. Rev. Microbiol. 63, 523–539 (2009).

21. Landa, M., Cottrell, M. T., Kirchman, D. L., Blain, S. & Obernosterer, I. Changes in bacterial diversity in response to dissolved organic matter supply in a continuous culture experiment. Aquat. Microb. Ecol. 69, 157–168 (2013).

22. Waterbury, J. B. & Valois, F. W. Resistance to co-occurring phages enables marine synechococcus communities to coexist with cyanophages abundant in seawater. Appl. Environ. Microbiol. 59, 3393– 3399 (1993).

23. Sullivan, M. B., Waterbury, J. B. & Chisholm, S. W. Cyanophages infecting the oceanic cyanobacterium Prochlorococcus. Nature 424, 1047–1051 (2003).

24. Suttle, C. A. Marine viruses--major players in the global ecosystem. Nat. Rev. Microbiol. 5, 801–812 (2007).

25. Thingstad, T. F. Elements of a theory for the mechanisms controlling abundance, diversity, and biogeochemical role of lytic bacterial viruses in aquatic systems. Limnol. Oceanogr. 45, 1320–1328 (2000)

26. Våge, S., Storesund, J. E. & Thingstad, T. F. SAR11 viruses and defensive host strains. Nature 499, E3–4 (2013).

27. Zhao, Y. et al. Abundant SAR11 viruses in the ocean. Nature 494, 357–360 (2013).

28. Stahl, D. A. & Torre, J. R. de la. Physiology and diversity of ammonia-oxidizing Archaea. Annu. Rev. Microbiol. 66, 83–101 (2012).

29. Timko, S. et al. Depth-dependent photodegradation of marine dissolved organic matter. Front. Mar. Sci. 2, doi:10.3389/fmars.2015.00066 (2015).

30. Noble, R. T. & Fuhrman, J. A. Virus decay and its causes in coastal waters. Appl. Environ. Microbiol. 63, 77–83 (1997).

## Supplementary References

31. Guillard, R. R. L. Culture of Phytoplankton for Feeding Marine Invertebrates. in Culture of Marine Invertebrate Animals: Proceedings — 1st Conference on Culture of Marine Invertebrate Animals Greenport (eds. Smith, W. L. & Chanley, M. H.) 29–60 (Springer US, 1975). doi:10.1007/978-1-4615-8714-9_3.

32. Caron, D. A. Enrichment, Isolation, and Culture of Free-Living Heterotrophic Flagellates. in Handbook of Methods in Aquatic Microbial Ecology (CRC Press, 1993), pp. 77–89.

33. Marie, D., Partensky, F., Vaulot, D. & Brussaard, C. Enumeration of phytoplankton, bacteria, and viruses in marine samples. Curr. Protoc. Cytom. Chapter 11, Unit 11.11 (2001).

34. Brussaard, C. P. D. Optimization of procedures for counting viruses by flow cytometry. Appl. Environ. Microbiol. 70, 1506–1513 (2004).

35. Rose, J. M., Caron, D. A., Sieracki, M. E. & Poulton, N. Counting heterotrophic nanoplanktonic protists in cultures and aquatic communities by flow cytometry. Aquat. Microb. Ecol. 34, 263–277 (2004).

36. Nadkarni, M. A., Martin, F. E., Jacques, N. A. & Hunter, N. Determination of bacterial load by real-time PCR using a broad-range (universal) probe and primers set. Microbiol. Read. Engl. 148, 257–266 (2002).

37. Sekiguchi, H., Watanabe, M., Nakahara, T., Xu, B. & Uchiyama, H. Succession of Bacterial Community Structure along the Changjiang River Determined by Denaturing Gradient Gel Electrophoresis and Clone Library Analysis. Appl. Environ. Microbiol. 68, 5142–5150 (2002).

38. Parada, A. E., Needham, D. M. & Fuhrman, J. A. Every base matters: assessing small subunit rRNA primers for marine microbiomes with mock communities, time series and global field samples. Environ. Microbiol. 18, 1403–1414 (2016).

39. Anonymous. 16S Metagenomic Sequencing Library Preparation. https://support.illumina.com/downloads/16s_metagenomic_sequencing_library_preparation.html. (27 November 2013).

40. Caporaso, J. G. et al. QIIME allows analysis of high-throughput community sequencing data. Nat. Methods 7, 335–336 (2010).

41. Bolger, A. M., Lohse, M. & Usadel, B. Trimmomatic: a flexible trimmer for Illumina sequence data. Bioinformatics 30, 2114–2120 (2014).

42. Zhang, J., Kobert, K., Flouri, T. & Stamatakis, A. PEAR: a fast and accurate Illumina Paired-End read merger. Bioinformatics 30, 614–620 (2014).

43. Callahan, B. J. et al. DADA2: High resolution sample inference from Illumina amplicon data. Nat. Methods 13, 581–583 (2016).

44. Quast, C. et al. The SILVA ribosomal RNA gene database project: improved data processing and web-based tools. Nucleic Acids Res. 41, D590–D596 (2013).

45. R Core Team. R: A language and environment for statistical computing. R Foundation for Statistical Computing, Vienna, Austria. URL https://www.R-project.org/ (2020).

46. Wickham H. ggplot2: Elegant Graphics for Data Analysis. Springer-Verlag New York. ISBN 978-3-319-24277-4, https://ggplot2.tidyverse.org (2016).

47. Csardi, G. & Nepusz, T. The igraph software package for complex network research. InterJournal, Complex Systems, 1695 (2006).

48. Aphalo, P. J. Learn R …as you learnt your mother tongue. Leanpub, Helsinki (2016).

